# Preferential associations of soil fungal taxa under mixed compositions of eastern American tree species

**DOI:** 10.1101/2021.08.09.455698

**Authors:** Steve Kutos, Elle M. Barnes, Arnav Bhutada, J.D. Lewis

## Abstract

Soil fungi are vital to forest ecosystem functions, in part through their role mediating tree responses to environmental factors, as well as directly through effects on resource cycling. While the distribution of these key taxa may vary with a suite of abiotic and biotic factors, the relative role of host species identity on soil fungal community composition and function remains unresolved. In this study, we used a combination of amplicon sequencing and enzymatic assays to assess soil fungal composition and associated function under three tree species, *Quercus rubra, Betula nigra*, and *Acer rubrum*, planted individually and in all combinations in a greenhouse, with added fungal inoculum collected below mature field trees. Across treatments, fungal communities were dominated by the phylum Ascomycota, followed by Basidiomycota and Mortierellomycota. Nonetheless, fungal communities differed between each of the solo planted treatments, suggesting at least some taxa may associate preferentially with these tree species. Additionally, fungal community composition under mixed sapling treatments broadly differed from solo saplings. The data also suggests that there were larger enzymatic activities in the solo treatments as compared with all mixed treatments. This difference may be due to the greater relative abundance of saprobic taxa found in the solo treatments. This study provides evidence of the importance of tree identity on soil microbial communities and functional changes to forest soils.

## INTRODUCTION

Forests provide globally important ecological and socio-economic functions (Pearce 2001, Butler-Leopold et al. 2018, Moser et al. 2020). They influence nutrient cycles, contain high biodiversity, and are central components of many terrestrial environments in terms of biomass, primary productivity, and land coverage (Pearce 2001, Moser et al. 2020). However, forest trees are under increasing stress from a multitude of factors including climate change, land-use change, habitat fragmentation, and invasive species (Dale et al. 2001, Butler-Leopold et al. 2018, Jo et al. 2019, Moser et al. 2020). Studies have demonstrated that soil fungi can mediate tree responses to these factors (van der Heijden et al. 1998, Hart et al. 2003, Smith & Read 2010, Fontaine et al. 2011, Karst et al. 2015). This interaction is highly complex: studies of the relative roles of trees and fungi in regulating these interactions have yielded differing results (De Deyn & Van der Putten 2005, Bardgett & van der Putten 2014, Lladó et al. 2018). For instance, plant diversity (Bardgett & van der Putten 2014, Moeller et al. 2015, Prober et al. 2015, Leff et al. 2018, Tedersoo et al. 2020), soil fungal diversity (van der Heijden et al. 1998, 2008, Mangan et al. 2010, Wagg et al. 2011), and soil characteristics (Zobel & Öpik 2014, Zhou et al. 2016, Baldrian 2017, Tedersoo et al. 2020) all have been suggested to be the principal factor(s) driving the functionality of this ecosystem dynamic. Therefore, it is critical for the conservation and management of forests that we improve our understanding of this complex tree-fungal interaction.

Soil fungi are globally distributed and are found in high relative abundances in forests (Rousk et al. 2010, Tedersoo et al. 2012, 2014, Talbot et al. 2014). These taxa provide essential ecosystem services including decomposition of organic matter and increasing tree access to limiting nutrients (van der Heijden et al. 1998, Lynd et al. 2002, Smith & Read 2010). It is well established that both abiotic and biotic factors can lead to spatial differences in soil fungal community composition (Martiny et al. 2006, Tedersoo et al. 2014, 2020). Key abiotic factors include climate characteristics such as precipitation, as well as soil characteristics such as pH and nutrient availability (Bardgett & Wardle 2010, Rousk et al. 2010, Tedersoo et al. 2012, 2020, Essene et al. 2017). Biotic factors influencing fungal communities include forest successional stage, microbial species pool, fungal taxa dispersal capabilities, and the aboveground tree diversity (Molina & Trappe 1982, Ishida et al. 2007, Buée et al. 2009, Turner et al. 2009, Bardgett & Wardle 2010, Peay & Bruns 2014). A central question, however, is the relative importance of each of these factors on fungal community composition (Guerra et al. 2020).

An unresolved component of this question is the role of tree distributions on the distribution of soil fungal communities. On the one hand, studies have shown a minimal influence of tree species distribution on soil fungal community composition, instead suggesting a primary influence of aforementioned characteristics (Ryberg et al. 2009, Smith et al. 2011, Kennedy et al. 2012, Botnen et al. 2014). Other studies have more closely tied belowground fungal distribution to tree identity and tree phylogenetic relatedness (Molina & Trappe 1982, De Deyn & Van Der Putten 2005, Ishida et al. 2007, Tedersoo et al. 2010, Lang et al. 2011, Molina & Horton 2015, van der Linde et al. 2018). While many soil-fungal taxa are generalists that associate with many tree species, certain taxa appear to have a narrow tree species preference, possibly due to their coevolution (Molina & Trappe 1982, Massicotte et al. 1999, Klironomos 2000). Fungal preferential association may also arise through several mechanisms, including preferential allotment of tree-derived photosynthate, microbial competition, and tree physiological differences (e.g., modification of soil micro-habitats through root exudates; Dickie 2007, Saunders & Kohn 2009, Aponte et al. 2010, Kiers et al. 2011). The role of fungal host preferential association may become more complicated when trees are in close proximity, leading fungal community composition to become mixed or possibly skewed towards the community associated with one tree species (Hubert & Gehring 2008, Hausmann & Hawkes 2009, Bogar & Kennedy 2013). A better understanding of the role of neighboring trees on soil fungal communities can provide further insights into how these communities assemble through time and space.

One way to uncover how the distribution of trees influences the distribution and function of soil fungi is understanding the relationship between fungal community composition and changes in fungal exudates. Extracellular enzymes released by fungi break down organic molecules in the local soil area, contributing to nutrient cycling (Burns 1982, Sinsabaugh & Moorhead 1994, Chalot & Brun 1998, Lynd et al. 2002, Burns et al. 2013). Functional redundancy among these taxa might be high, nonetheless a change in fungal composition may alter enzymatic activities in a local area, altering decomposition rates and soil nutrient storage or release (Burns et al. 2013, Kyaschenko et al. 2017, Looby & Treseder 2018, Nannipieri et al. 2018). Therefore, spatial differences in fungal community composition associated with differences in tree distributions could lead to altered enzymatic activity, which in turn could impact fungal community composition through effects on nutrient cycling. Accordingly, it is critical to assess changes in production of key enzymes associated with carbon flux and nutrient cycling, and their effects on relationships between tree and soil fungal distributions.

For this study, we examined if soil fungal communities associated with three tree species are distinct from one another and mix if the trees are in close proximity. To address this, we used a greenhouse bioassay with saplings of three common eastern USA tree species, *Quercus rubra, Betula nigra*, and *Acer rubrum*, planted individually as well as in paired and triplicate combinations. A fungal inoculate was added to these microcosms, which was collected from soils below mature trees of these same species in the field. We hypothesized that the soil fungal communities of the three tree species would be distinct from one another and decrease in the mixed treatments, due to changes in niche availability and/or interspecific competition. We also examined how potential differences in fungal community composition might result in functional changes in the form of potential activity for six common fungal extracellular enzymes. Finally, we hypothesized that soil enzymatic activity would differ across treatments and will be significantly greater in treatments containing a higher relative abundance of saprobic functional taxa. While interpreting the results of greenhouse bioassays is limited by their simplicity compared to actual forest ecosystems, such studies also minimize confounding effects of abiotic factors that might obscure interpretation of the effects of tree species and fungal community function.

## METHODS

### Experimental bioassay design

To explore our hypothesies, we performed a greenhouse bioassay at Fordham University’s Louis Calder Center in Armonk, NY, USA (coordinates: 41.131789, -73.732911). The Calder Forest is a 113-acre protected preserve with a woodland comprising of oaks (*Quercus* spp.), maples (*Acer* spp.), American beech (*Fagus grandifolia*), hickory (*Carya* spp.), and birch (*Betula* spp.). The climate is temperate with mean annual temperatures of 12°C and mean annual precipitation of 120 cm. The soil is acidic sandy loam. Experimental tree species were obtained as ∼30 cm tall bareroot saplings from Cold Stream Farm (Freesoil, MI, USA). In autumn 2018, each sapling was planted individually in a sterilized 3.8 cm-wide cone-tainer pot (Steuwe and Sons, Tangent, OR, USA) with field-collected soils. These soils were collected from underneath five randomly selected mature trees for each experimental species at least 30 m apart in the Calder Forest. Roughly 6 kg of soil for each tree species was collected and homogenized so that each experimental sapling received the same soil source.

In early spring 2019, saplings of similar height of each species were randomly selected from this stock for use within the bioassay. Saplings were then moved into sterilized C1200 11-liter plastic pots with a 2:1 volumetric mixture of soil mix (*Pro-Mix BX*, Premier Tech Horticulture, Quakertown, PA, USA) and autoclaved sand (Sakrete, Charlotte, NC, USA). *Pro-Mix BX* soil is a general-use, non-mycorrhizal mixture containing sphagnum peat moss, perlite, limestone, and vermiculite. In the greenhouse, the pots were arranged in a randomized block design with ∼15 cm spacing between pots to reduce crowding (12 blocks x 8 treatments). Treatments were: (1) *Controls:* pots with no trees, (2) *Solo Planted:* one sapling from each species planted alone, (3) *Paired Planted:* two different tree species planted together in all pairwise combinations, and (4) *Triple Planted:* all tree species planted together. For the pair and triple plantings, each individual was placed in the pot roughly equidistant apart with a few centimeters spacing from the edge. To these containers, a soil fungal inoculum was added for each species within the treatment to obtain a representative fungal diversity mimicking field condition. The fungal inoculate were created from soils taken from underneath the same trees used for the initial planting substrate and contained a mixture of 500 mL sterile diH_2_O and 100 mL soil. Controls received the same 2:1 ratio of soil to autoclaved sand, but without added inoculum, to account for fungal taxa within the *Pro-Mix BX*. Soil moisture and pH were evaluated monthly to confirm treatments were under similar conditions during the experiment. Soil pH was measured using a 2:1 ratio of distilled water to soil on an Accumet AE150 probe (ThermoFisher Scientific, Waltham, MA, USA), and soil moisture was measured by drying 5 g of soil for 24 h at 105° C.

### Soil sampling, DNA sequencing, and bioinformatics

In early winter 2019, three soil cores (∼8 cm depth) were collected from each pot and pooled into one soil sample per treatment. The soil corer was sterilized with 10% bleach and 70% ethanol prior to each sampling. Soil cores were sieved through a sterilized 2 mm screen and stored at -20° C. Fungal DNA was extracted from 0.25 g of soil using the Qiagen PowerSoil DNA Kit (Qiagen Inc., Mississauga, Canada) and PCR targeted the ITS1 region using the forward primer ITS1F and barcoded reverse primer ITS2 (White et al. 1990, Gardes & Bruns 1993, Smith & Peay 2014). PCR reactions (25 μl) contained: 0.25 μl Platinum *Taq* (5 U/μl), 2.5 μl 10X Invitrogen Buffer, 0.75 μl 50 mM MgCl_2_, 1.0 μl of both primers (10μM), 0.5 μl 10mM dNTPs, 1.0 μl BSA, and 2 μl of extracted DNA. Amplifications were performed on an Applied Biosystems thermocycler (Model 2720, Foster City, CA, USA) under the following conditions: 94° C for 1 min, followed by 94° C for 30 s, 58° C for 30 s, 68° C for 30 s for 35 cycles, and 68° C for 7 min. Amplicons were purified using Sera-Mag Speedbeads™ (GE Healthcare, Chicago, IL, USA; Rohland & Reich 2012) and quantified with a Qubit 4.0 Fluorometer (ThermoFisher, Carlsbad, CA, USA). Amplicon libraries, including positive and negative controls, were normalized, pooled, and then sequenced on an Illumina 2 × 250 bp MiSeq at Genewiz (Brooks Life Sciences Company, South Plainfield, NJ, USA). The positive sequencing control contained ten known fungal taxa from the phyla Basidiomycota, Ascomycota, and Mortierellomycota. Demultiplexed FASTQ files were filtered and processed using a QIIME2 bioinformatic pipeline (Release: 2020.8; Caporaso et al. 2010). Sequencing adapters, fungal primers, and low-quality bases were first trimmed using *ITSXpress* and *cutadapt* (Martin 2011, Rivers et al. 2018). Trimmed reads were processed and merged using DADA2 to produce ASVs (Callahan et al. 2016) and then aligned via the UNITE database to assign them to taxonomic groups (Version 8.2, Abarenkov et al. 2010). ASVs that did not match taxa within the database were listed as unclassified fungal ASVs. ASVs classified as non-fungi were removed. These ASVs were rarified to a depth of 7025. Finally, taxonomic assignments were processed in *FUNGuild* to assign ASVs to functional groups using only highly probable and probable assignments (Nguyen et al. 2016).

### Fluorometric enzymatic assay

To obtain fungal enzymatic potential activity, soils were run through a high-throughput standardized fluorescence assay developed by Bell *et al*. (2013) using 4-methylumbelliferone as a fluorescent indicator (Jacks & Kircher 1967, Marx et al. 2001, Courty et al. 2005). Six evaluated enzymes were selected due to their abundance in soils and their role in the degradation of carbon and phosphorus substrates (**Supplemental Table 1**; Nannipieri et al. 2012, Burns et al. 2013). Factorial tests runs were conducted to determine maximum yields using different pH levels found in the area (4.0 – 6.0), addition of NaOH, different incubation times (3 – 6 hours), and multiple read settings as recommended in the protocol (Bell et al. 2013). Soil slurries for each sample were created by combining 2.75 g of soil and 91 ml of 50 mM sodium acetate buffer. From these slurries, 800 ul of each sample was added randomly to a 96 deep-well plate and incubated for 3 h in the dark at room temperature (∼23° C) and then centrifuged for 30 min at 2300 x g. From the supernatant, 200 ul of each sample was added to corresponding black, flat-bottom plates and read on a SpectraMax M2e microplate reader (Molecular Devices, San Jose, CA, USA), with an excitation wavelength at 365 nm and emission wavelength at 450 nm. To improve fluorescence yields, 10 ul of NaOH was added to each well roughly two minutes prior to plate reading (DeForest 2009, Bell et al. 2013). All assays were completed within 72 h of soil collection to avoid suppressing activity (DeForest 2009). Negative controls consisting of only a solution of the buffer and substrate used for each sample. Enzymatic activities were corrected by applying this control and converted to nmol h^−1^ g^−1^.

### Statistical analysis

All analyses were completed in R (Version 4.0.1) using the *phyloseq, vegan, microbiome* packages (Oksanen et al. 2007, McMurdie & Holmes 2013, Lahti et al. 2017). Alpha diversity was measured as ASV richness and by the Shannon index and diversity was compared among treatments using an ANOVA including block as a random effect, followed by Tukey’s HSD tests for pairwise comparisons. Levene’s tests and histograms were used to assess normality and homogeneity of variances. Beta diversity was measured by calculating Bray-Curtis dissimilarities on log-transformed read counts. Homogeneity of dispersions was checked using *betadisper*, and PERMANOVAs were run to test for significance using *adonis* and visualized using an NMDS ordination. Pairwise analysis of beta diversity differences among locations was measured using the function *pairwise*.*adonis* with an FDR correction for multiple comparisons (Martinez Arbizu 2020). Relative abundance heatmaps were created on the 20 most abundant ASVs in each of the major phyla using the function *aheatmap* with the rows converted to standardized Z-Scores. Indicator species analysis was performed to examine which ASVs were distinctive to each treatment using the *indicspecies* package with 9999 Monte Carlo permutations (Cáceres & Legendre 2009). Enzyme values obtained from the fluorometric assay were compared using an ANOVA with a Tukey’s HSD for pairwise comparisons. Enzymatic activity values underwent a log transformation prior to analysis due to skewness. Graphs were constructed using the ggplot2 package (Wickham et al. 2016).

## RESULTS

### Soil fungal community diversity patterns

In total, our samples contained 1,581 unique ASVs (3.65 × 10^5^ sequences from 52 samples). The phylum with the greatest number of ASVs across all treatments was Ascomycota (82.9% of all reads), with lesser numbers in the Basidiomycota (14.0%), and Mortierellomycota (2.5%; **Fig. 1**,**2**). The ten most relative abundant ASVs across all treatments were also members of Ascomycota (orders: Pezizales, Xylariales, Saccharomycetales, Coniochaetales, Thelebolales) and Basidiomycota (order: Tremellales). There were only two ASVs found within all treatments belonging to the order Saccharomycetales within Ascomycota.

**Figure 1.**
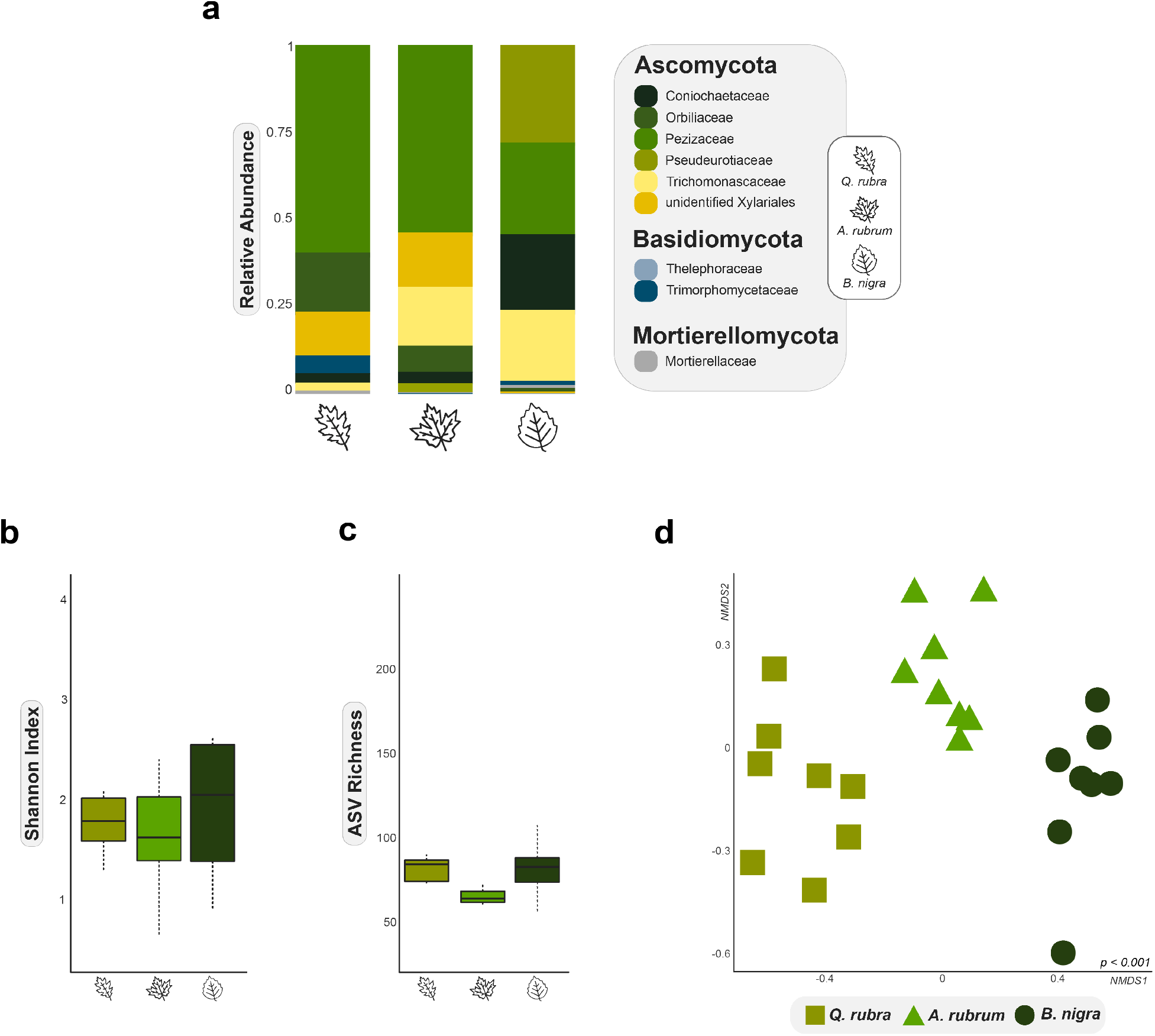
Soil Fungal Diversity in Solo Planted Treatments. a: Relative abundance of the top nine families for all solo treatments. b: Boxplot of Shannon index of soil fungal communities among solo treatments. c: Boxplot of ASV richness of soil fungal communities among solo treatments. d: NMDS ordination of Bray-Curtis distances of the solo treatments.

**Figure 2.**
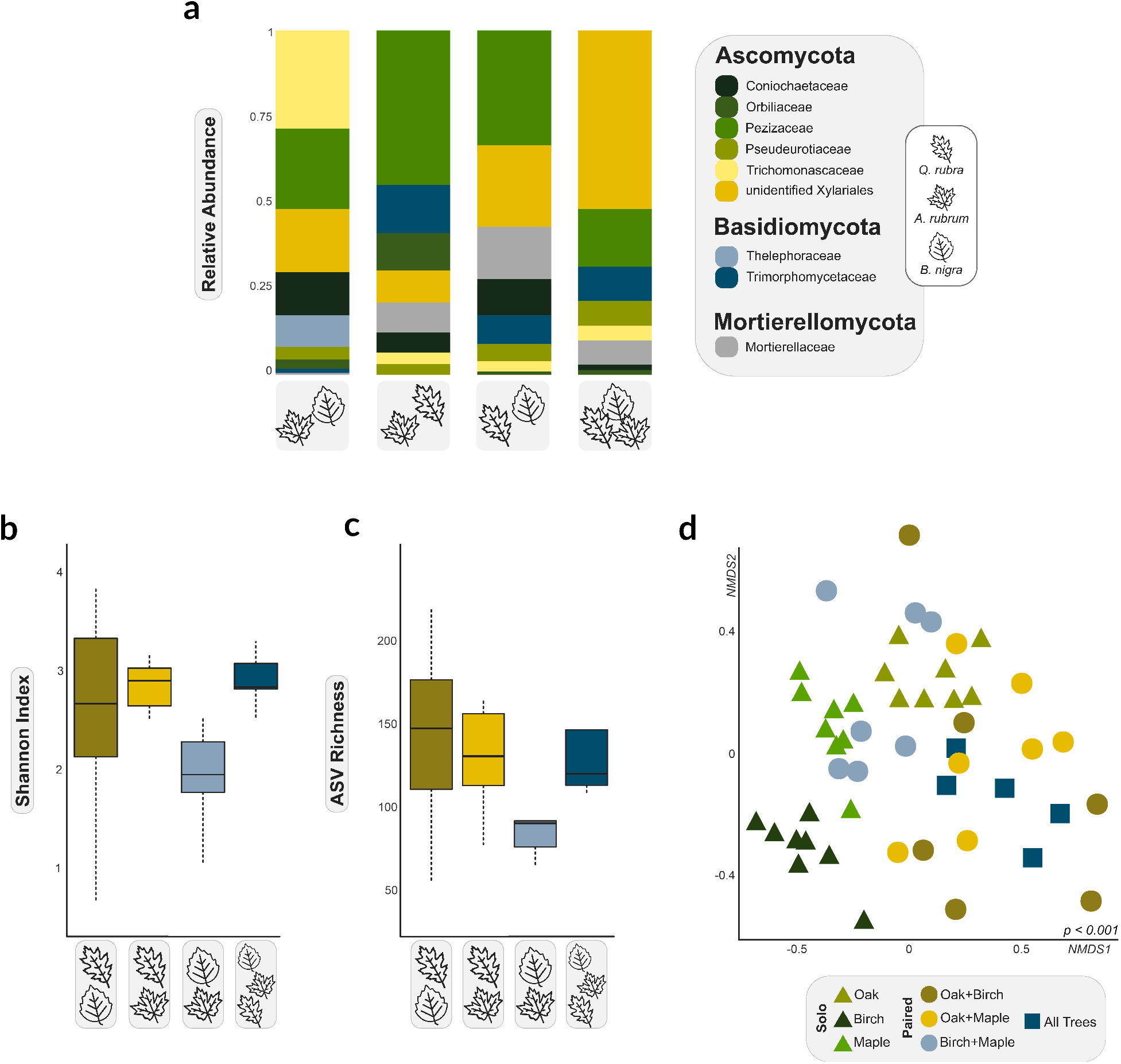
Soil Fungal Diversity in Mixed Planted Treatments. a: Relative abundance of the top nine families for all mixed treatments. b: Boxplot of Shannon index of soil fungal communities among mixed treatments. c: Boxplot of ASV richness of soil fungal communities among mixed treatments. d: NMDS ordination of beta diversity of all solo and mixed treatments.

All solo planted treatments had a greater relative abundance of the phylum Ascomycota as compared to the mixed planted treatments (**Fig. 1, 2, 3a**). Alongside this shift, there were also clear differences in the relative abundance of the other two fungal phyla, Basidiomycota or Mortierellomycota, from the solo treatments to the mixed planted treatments. The solo *Quercus rubra* treatments had the greatest relative abundance of the fungal family Pezizaceae (53% of reads), along with the family Orbiliaceae (14.9%), and an unidentified family within the order Xylariales (11.3%; **Fig. 1a**). The solo *Acer rubrum* treatments also had the greatest relative abundance of Pezizaceae (50.4% of reads), along with an unidentified family within the order Xylariales (15.7%), as well as the family Coniochaetaceae (14.5%, **Fig. 1a**). In the solo *Betula nigra* treatments, the families Pezizaceae (18.5% of reads), Pseudeurotiaceae (15.4%), Coniochaetaceae (14.5%, **Fig. 1a**) had the highest relative abundance. ASVs in Pezizaceae in the solo *B. nigra* treatments were ∼2.5X lower in relative abundance from the other solo treatments.

**Figure 3.**
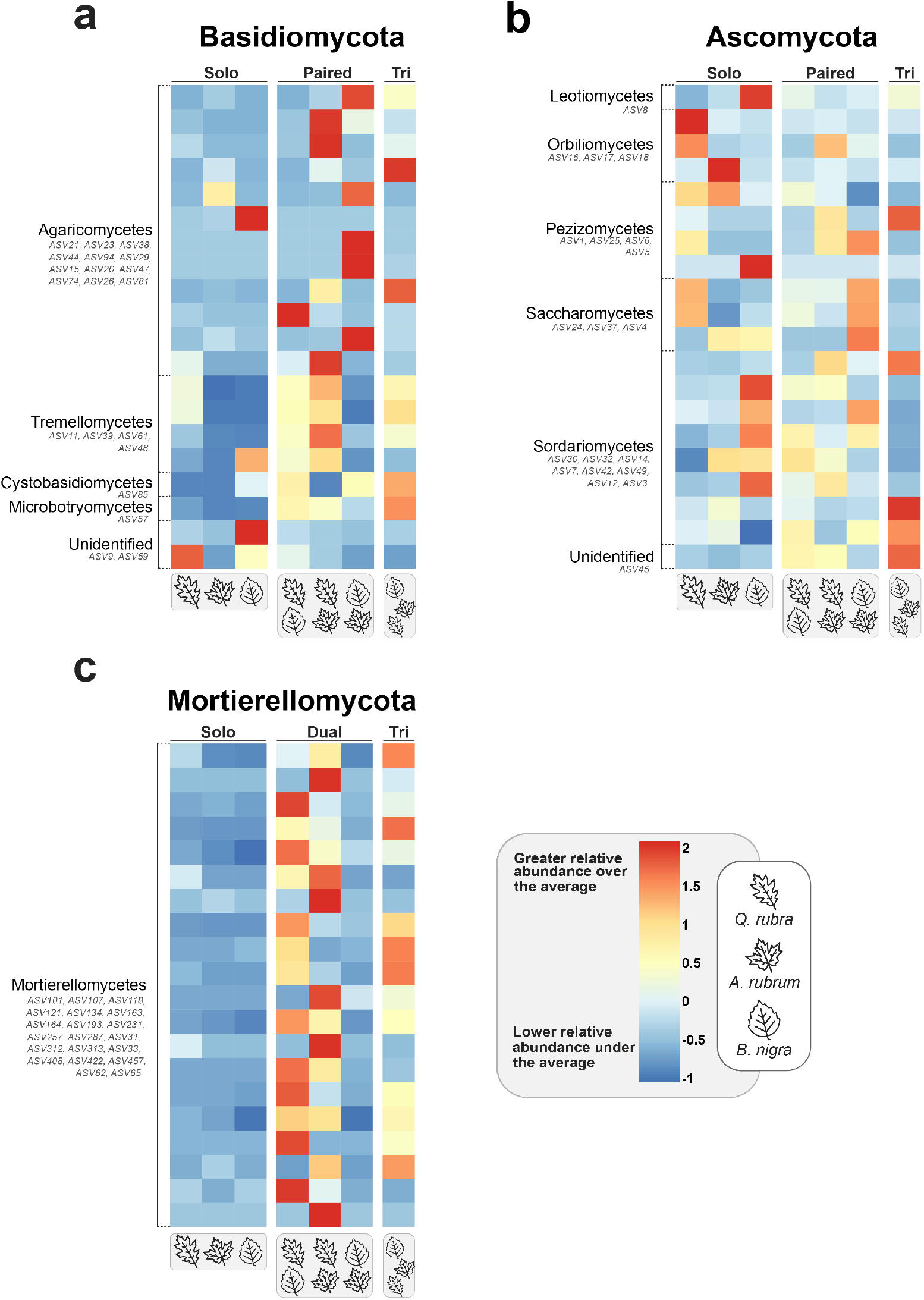
Distribution of fungal ASVs across treatments using *Z-Scores*. a: 20 most abundant Basidiomycota ASVs across treatments. b: 20 most abundant Ascomycota ASVs across treatments. c: 20 most abundant Mortierellomycota ASVs across treatments. Fungal classes are listed on the left of each graph.

When *Q. rubra* was paired with *A. rubrum*, taxa in the family Pezizaceae (32% of reads) and the unidentified family within the order Xylariales (6.3%) had lower relative abundances by 1.6X and 1.8X respectively compared to the solo *Q. rubra* treatments (**Fig. 2a**). However, these two families had the highest relative abundance in this paired treatment along with the family Trimorphomycetaceae (8.4%, **Fig. 2a**). Taxa in the family Trichomonascaceae, which were highly abundant in the solo *A. rubrum* treatments, were 7X lower in abundance when paired with *Q. rubra*, matching the relative abundance in the solo *Q. rubra* treatments (**Fig. 2a**). Further, taxa within Mortierellaceae were 6X greater in relative abundance in this paired treatment when it was rare in both solo sapling treatments (< 0.1% of reads; **Fig. 2a, 3c**). When *B. nigra* was paired with *A. rubrum*, the diversity pattern changed with taxa in the family Trichomonascaceae (26.4% of reads) having the highest relative abundance along with an unidentified family in the order Xylariales (16.2%) and the family Pezizaceae (16.0%; **Fig. 2a**). Therefore, only in this paired treatment does the family Trichomonascaceae have a higher relative abundance as compared to both solo sapling treatments. The family Pseudeurotiaceae, which was highly abundant in the solo *B. nigra* treatments, had a 5X lower relative abundance, similar to abundances found in the solo *A. rubrum* treatments (2.8% of reads; **Fig. 2a**).

When *B. nigra* was paired with *Q. rubra*, the families Pezizaceae (26.7% of reads), an unidentified family in the order Xylariales (18.8%), and Mortierellaceae (7.9%) had the highest relative abundances (**Fig. 2a**). Thus, taxa within Mortierellaceae were 6X greater in relative abundance only in this paired treatment compared the solo sapling treatments (**Fig. 2, 3c**). Highly abundant families in the solo *B. nigra* treatments were lower in relative abundance only when paired with *Q. rubra*, including Pseudeurotiaceae (15.4% to 3.8% of reads), Coniochaetaceae (14.5% to 7% of reads), and Trichomonascaceae (13.7% to 2% of reads). This pattern was also seen with high relative abundant families in the solo *Q. rubra* treatments including Orbiliaceae (15% to 0.8% of reads) and Pezizaceae (53% to 27% of reads).

When all three tree species were planted together, the highest relative abundance was in the taxa in an unidentified family in the order Xylariales (35.5%), Pezizaceae (11.3%), and Trimorphomycetaceae (6.3%; **Fig. 2**). Taxa in Trichomonascaceae and Coniochaetaceae, which had high relative abundances in the *B. nigra* and *A. rubrum* solo treatments, were 4X and 10X less abundant with respect to the triple-plated treatment (**Fig. 2**). While only found in small relative abundances, ectomycorrhizal fungal taxa were 6X higher in the paired and triple sapling treatments as compared with the solo treatments, whereas saprobic taxa displayed no distinct pattern.

Fungal communities differed significantly among treatments in Shannon diversity (*F* = 3.78, *p* = 0.003; **Fig. 1b, 2b**) and ASV richness (*F* = 7.49, *p* < 0.001; **Fig. 1c, 2c**). Diversity and ASV richness in solo treatments differed from that in the mixed sapling treatments, with greater diversity (∼1.5X) and richness (∼1.5X) in the paired and triple-plated treatment (**Fig. 1, 2**). Community composition also differed significantly (*F* = 3.35, *p* < 0.001, r^2^ = 0.22; **Fig. 1, 2**). There were significant differences in the fungal composition between each solo planted treatment as well as between the solo planted treatments and mixed planted treatments (*p* < 0.01; **Supplemental Table 2**). When we looked only at the mixed planting treatments, we found significant differences in the fungal composition of all paired treatments except between the *Q. rubra + A. rubrum* and the *Q. rubra + B. nigra* treatments (*p* < 0.01; **Supplemental Table 2**). Finally, we found no differences in composition between the triple-planted and paired-planted treatments except for the triple-planted versus *B. nigra* + *A. rubrum* treatment (*p* < 0.01; **Supplemental Table 2**).

Indicator species analysis was used to detect ASVs that may significantly associate with one tree species over the others or with one tree species combination. Five fungal ASVs were identified as indicator ASVs in the solo- and paired planted treatments (*p* < 0.01). Of note, the relative abundances of these indicator taxa were low (< 0.1% of reads). An unidentified ASV in the family Herpotrichiellaceae and an ASV in the family Geminibasidiaceae were identified as indicator ASVs in the solo *Q. rubra* treatments. The solo *A. rubrum* treatments contained one indicator ASV in the order Glomerales. The solo *B. nigra* treatments also had one indicator ASV in the family Mycosphaerellaceae. The paired treatment of the *Q. rubra* + *A. rubrum* had one indicator ASV in the family Trimorphomycetaceae. The remaining treatments included no indicator ASVs (*p* > 0.01).

### Enzyme activity patterns

Two enzymes assessed, α-Glucosidase and β-glucuronidase, displayed no detectable activity via the fluorometric assay, so they were removed from statistical analyses. The activities for the remaining four enzymes were significantly different among treatments except for β-xylosidase, which did not vary by treatment (*p* > 0.05; **Fig. 4c**). The enzymatic potential activity of β-Glucosidase was significantly different between the treatments. This difference was mainly detected in the solo *A. rubrum* and solo *B. nigra* treatments, which had β-Glucosidase activity levels that were 3X higher than all other treatments (*p* < 0.001; **Fig. 4a**). The enzymatic potential activity of cellobiohydrolase was also significantly different between treatments. In particular, cellobiohydrolase activity was 4X higher in the solo *A. rubrum* and *B. nigra* treatments (**Fig. 4d**). Acid phosphatase activity was 3.5X higher in all three solo treatments as compared to all mixed treatments (*p* < 0.001; **Fig. 4b**). As expected, the no-sapling controls showed no discernible activity for any of the enzymes evaluated.

**Figure 4.**
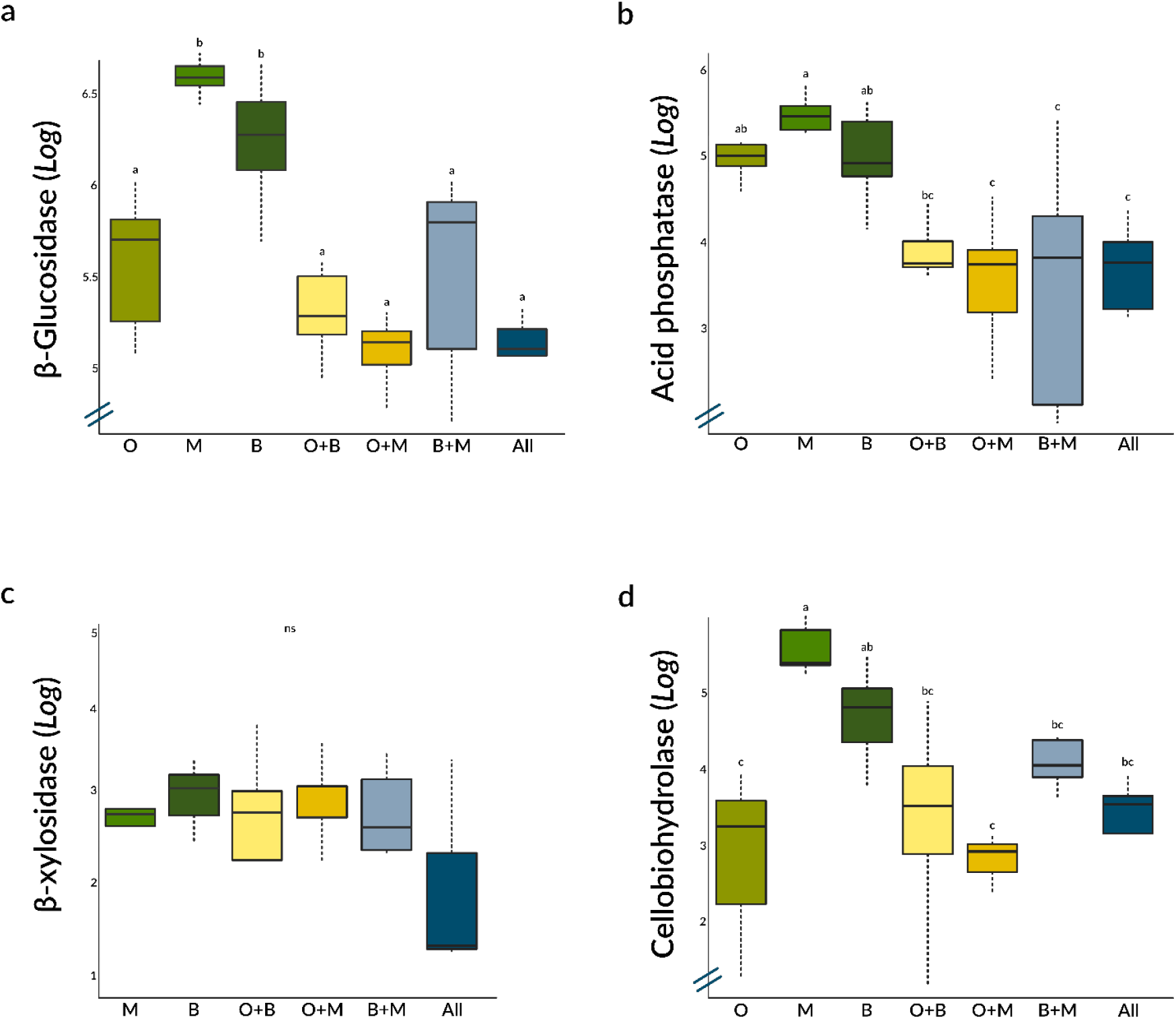
a: Boxplot of log-transformed enzymatic activity for β-glucosidase (*p* < 0.05). b: Boxplot of log-transformed enzymatic activity for acid phosphatase (*p* < 0.05). c: Boxplot of log-transformed enzymatic activity for β-xylosidase (ns, *p* > 0.05). d: Boxplot of log-transformed enzymatic activity for cellobiohydrolase (*p* < 0.05). **O** = solo *Q. rubra*, **M** = solo *A. rubrum*, **B** = solo *B. nigra*, **O+B** = paired *Q. rubra* and *B. nigra*, **O+M** = Paired *Q. rubra* and *A. rubrum*, **B+M** = paired *B. nigra* and *A. rubrum*, and **All** = treatment of all three experimental species. All values are nmol h^−1^ g^−1^. Letters denote significant Tukey’s HSD.

## DISCUSSION

This study investigated the effects of tree identity and composition on fungal communities in temperate forest soils. Overall, soil fungal community composition differed with tree sapling species identity. Further, the soil fungal communities of the solo planted treatments differed from the mixed planted treatments, indicating that these three tree species can develop distinct fungal communities and that when proximal, their associated fungal communities’ mix. In addition, we observed that differences in fungal community composition were associated with altered activity of enzymes. By reducing environmental variation associated with field studies, this greenhouse bioassay provides a more direct exploration of tree-host preferential association among soil fungal taxa.

Consistent with our first hypothesis regarding solo planted treatments, the soil fungal communities associated with each of the three tree species, *Acer rubrum, Quercus rubra*, and *Betula nigra*, differed in community diversity and composition. This was anticipated due to the relatively distant phylogenetic relatedness of these tree species, which has a positive relationship with soil fungal community composition (Molina & Trappe 1982, Ishida et al. 2007, Tedersoo et al. 2013). Overall, there were taxonomic patterns distinguishing these communities. As one example, taxa in the fungal class Leotiomycetes (phylum: Ascomycota) and fungal families Pyronemataceae (phylum: Ascomycota) and Phanerochaetaceae (phylum: Basidiomycota) were only highly abundant in the solo *B. nigra* treatments suggesting robust tree species preferential association. These fungal groups do have known associations with *Betula* spp., although not exclusively, suggesting potential preferential association, but not specificity (Tedersoo & Smith 2013, Hrynkiewicz et al. 2015, Miettinen et al. 2016, Ekanayaka et al. 2019). These findings provide further evidence that tree species can associate with distinct fungal communities.

When we compared both solo and mixed planted treatments, we found a few fungal families that had high relative abundances in all treatments. This implies that most of the fungal taxa found within these communities are tree species or environmental condition generalists and might be able to associate with multiple tree species (Ishida et al. 2007, Urbanová et al. 2015). This is further bolstered by the lack of an broad influence of one tree species’ distinct fungal community ‘overwhelming’ the communities of the other trees (Mummey et al. 2005, Hausmann & Hawkes 2009). However, soil fungal community composition in the solo planted treatments was significantly different from the mixed planted treatments. For instance, taxa in the fungal family Tremellomycetes (phylum: Basidiomycota) were only abundant in treatments containing a *Q. rubra*. These taxa are common soil residents in temperate forests and have diverse morphologies and nutrient acquisition strategies (Edwards & Zak 2010, Millanes et al. 2011, Liu et al. 2015, Mašínová et al. 2017). Their apparently specific association in the present study with *Q. rubra* is of interest and warrants further study. A different pattern was found with the family Trichomonascaceae (phylum: Ascomycota) which was only abundant in treatments lacking *Q. rubra*, suggesting either preferential association for *B. nigra* and *A. rubrum*, or a preferential aversion to *Q. rubra*. This family of yeasts has seen limited study; however, they do belong to a fungal group that may be sensitive to increased microbial competition and changes in nutrient conditions (Kurtzman & Robnett 2007, Li et al. 2021).

Greater fungal richness and diversity in the mixed planted treatments seemed to be driven at least partially by the higher relative abundances of taxa from the phyla Mortierellomycota and Basidiomycota. Taxa within Mortierellomycota are common soil saprobic residents that can form a mycorrhizal-like symbiosis with many tree hosts (Uehling et al. 2017, Johnson et al. 2019). Similarly, some taxa within Basidiomycota prefer colonizing the roots of multiple hosts (Molina & Trappe 1982, Horton & Bruns 1998, Massicotte et al. 1999, Smith & Read 2010). This may help to explain why these taxa had higher relative abundances in the mixed planted treatments, especially given that saplings in the bioassay were planted relatively close together. Further, studies have suggested that when certain fungal taxa are already colonized on one tree root, proximal plants can prompt sporulation and germination, thus increasing the opportunity of multi-tree colonization (Hubert & Gehring 2008, Bogar & Kennedy 2013, Bogar et al. 2015). Interestingly, Basidiomycota taxa had different distributions between the mixed treatments depending on tree composition. For example, closely related genera *Lactarius* and *Russula* (order: Russulales) were more abundant in separate mixed treatments (*Lactarius: Q. rubra + A. rubrum* and triple-planted treatments; *Russula: Q. rubra + B. nigra* treatments). This pattern further supports evidence from other studies that these genera can be host-specific if preferred hosts are available but also host generalists if only non-preferred hosts are available (Tedersoo et al. 2008, Morris et al. 2009, Wilson et al. 2017, Looney et al. 2018). In sum, our results suggest that the mixed plantings may work to synergistically provide a more suitable environment for certain fungal taxa, leading to their relative abundances being greater.

Taxa within the fungal phylum Ascomycota had markedly higher relative abundance in this bioassay compared to field studies in the region (Ramirez et al. 2011, Reese et al. 2016, Barnes et al. 2021), which could be due to the inoculum used and/or the greenhouse setting. These taxa are known to disperse easily by spore and hyphal fragments, can colonize multiple niches, can utilize multiple resources, have high-stress tolerance, and may represent generalists adaptable to greenhouse settings (Yamamoto et al. 2012, Prober et al. 2015, Egidi et al. 2019). This may explain why there was a high relative abundance of the genera *Chromelosporium* and *Sphaerosporella* which are commonly found within greenhouse experiments (Hennebert & Korf 1975, Sánchez et al. 2014).

Finally, it is important to acknowledge that there was a considerable amount of unexplained compositional variation within these treatments. This indicates that our bioassay communities are likely impacted by unmeasured factors, such as a legacy effect (Gao et al. 2020). Our mixed planted fungal inoculum consisted of a combination of soils found under each of the tree species in the field, rather than a single soil inoculum from a location where all three tree species grew side-by-side. We also cannot rule out the influence of artificially combining our fungal inoculum with any communities found in our greenhouse potting soil. Both events could have led our bioassay communities to mix in a slightly different way than might naturally happen in the field (Rillig et al. 2016, Castledine et al. 2020). Nonetheless, these findings might have important ecological impacts in eastern USA forests as fungal taxa that may be generalists, especially mycorrhizal fungi, could facilitate heterospecific sapling establishment and sapling development (Nara 2006, Dickie & Reich 2005, Dickie 2007, Wolfe & Pringle 2012).

### Enzymatic activity shifts among different tree compositions

Tree identity and composition have the potential to influence the enzymatic activity in soils by creating distribution patterns of fungal taxa with specific functional roles. The activities of β-glucosidase, cellobiohydrolase, and acid phosphatase were significantly higher in the solo planted treatments. One potential explanation for this is the higher relative abundance of particular saprobic taxa, which are known to emit extracellular enzymes, as well as the higher relative abundance of mycorrhizal taxa in the mixed plantings, which normally do not exhibit robust enzymatic activity (López et al. 2004, Kurtzman & Robnett 2007, Damm et al. 2010, Boberg et al. 2011, Péter et al. 2012). That the higher relative abundance of taxa in Mortierellomycota in the mixed treatments did not correlate with higher enzymatic activity is not surprising as while they are saprobic taxa, the main ASV found, *Mortierella elongata*, may be able to utilize simple sugars and not complex polymers (Poll et al. 2010, Ellegaard-Jensen et al. 2013, Uehling et al. 2017, Koechli et al. 2019).

Of note, these soils did not significantly differ in sapling age or soil characteristics such as pH and soil moisture levels which are known influencers of enzymatic activity differences (Brockett et al. 2012, Baldrian 2014, Kivlin & Treseder 2014). However, other untested soil conditions, as well as root exudates, may have contributed to the differences in enzymatic activities (Strickland et al. 2009, Kivlin & Treseder 2014). Additionally, only six enzymes were evaluated individually here which may miss other important enzymes present within these soils and/or the potential synergistic effect of multiple enzymes on soil substrates (Baldrian et al. 2014, Kivlin & Treseder 2014). Soil enzymatic activity is a dynamic process influenced by many factors, including other microbial groups, so caution should be used in interpreting this bioassay data (Boer et al. 2005, Romaní et al. 2006, Strickland et al. 2009, Glassman et al. 2018). Nevertheless, these results do add evidence to the relative importance of tree species and associated soil fungal communities on extracellular enzymatic activity.

## Conclusion

Data from this greenhouse bioassay indicate that different tree species can have significantly different soil fungal communities and that these communities can influence the fungal communities of proximal trees. These results match those from other tree-fungal systems. Additionally, the mixing of these fungal communities was associated with lower potential enzymatic activity. Forests in the eastern USA are not static; tree diversity and composition are changing alongside a changing climate. This data provides further evidence that altering the distribution and diversity of aboveground trees in these forests may impact belowground soil fungal communities. We acknowledge that this greenhouse bioassay was performed on young saplings which may have resulted in an increased influence of spore-based, early arriving fungal taxa. However, as this and other studies demonstrate, the diversity and richness patterns found here have the potential to relate to fungal communities *in situ*. To further explore these relationships, future experiments could include varied environmental conditions, more distant phylogenetic tree species, as well as a longer experimental period. Disentangling the factors that influence forest trees and soil fungal is of utmost importance given the vast array of ecosystem services provided by these two intertwined communities.

## Acknowledgments

We thank the Louis Calder Center for use of their facilities including the Calder Forest and the greenhouse. We also thank Drs. Steven Franks, John Wehr, and Patricio Meneses for their feedback on earlier drafts of this manuscript.

---

**Supplemental Table 1.** List of enzymes evaluated, the associated reagents used, and their ecological roles.

| Enzyme | Reagent | Enzymatic Action |
| --- | --- | --- |
| *Cellobiohydrolase (CBH)* | 4-MUB-β-D-cellobioside | **C Cycle:** Degrades cellulose |
| *β-Glucosidase (BG)* | 4-MUB β-D-glucopyranoside | **C Cycle:** Degrades cellulose |
| *α-Glucosidase (AG)* | 4-MUB α-D-glucopyranoside | **C Cycle:** Degrades cellulose |
| *Acid phosphatase (AP)* | 4-MUB Phosphate | **P cycle:** Phosphorus mineralization |
| *β-xylosidase (BX)* | 4-MUB β-D-xylopyranoside | **C Cycle:** Degrades cellulose & hemicellulose |
| *β-glucuronidase (BGLU)* | 4-MUB β-D-glucuronide | **C Cycle:** Degrades hemicellulose |

**Supplemental Table 2.** Pairwise comparisons of Bray-Curtis distance between all treatments. *Denotes significance with FDR correction.

| Comparison | Sum of Squares | F | r<sup>2</sup> | Adjusted p |
| --- | --- | --- | --- | --- |
| **Solo vs Solo** |  |  |  |  |
| Maple vs Birch | 0.675 | 4.585 | 0.247 | **0.001\*** |
| Maple vs Oak | 0.655 | 4.154 | 0.229 | **0.001\*** |
| Birch vs Oak | 1.080 | 6.550 | 0.319 | **0.003\*** |
| **Solo vs Paired** |  |  |  |  |
| Maple vs Birch & Maple | 0.517 | 3.087 | 0.192 | **0.001\*** |
| Maple vs Oak & Maple | 0.864 | 4.975 | 0.277 | **0.001\*** |
| Maple vs Oak & Birch | 0.820 | 4.308 | 0.264 | **0.001\*** |
| Birch vs Birch & Maple | 0.691 | 3.943 | 0.233 | **0.003\*** |
| Birch vs Oak & Maple | 0.928 | 5.106 | 0.282 | **0.001\*** |
| Birch vs Oak & Birch | 0.880 | 4.423 | 0.269 | **0.004\*** |
| Oak vs Birch & Maple | 0.468 | 2.511 | 0.162 | **0.001\*** |
| Oak vs Oak & Maple | 0.402 | 2.086 | 0.138 | **0.002\*** |
| Oak vs Oak & Birch | 0.438 | 2.072 | 0.147 | **0.007\*** |
| **Solo vs All** |  |  |  |  |
| Maple vs All Trees | 0.766 | 4.730 | 0.301 | **0.001\*** |
| Birch vs All Trees | 0.919 | 5.366 | 0.328 | **0.003\*** |
| Oak vs All Trees | 0.492 | 2.666 | 0.195 | **0.001\*** |
| **Paired vs Paired** |  |  |  |  |
| Birch & Maple vs Oak & Maple | 0.543 | 2.631 | 0.180 | **0.003\*** |
| Birch & Maple vs Oak & Birch | 0.542 | 2.382 | 0.178 | **0.001\*** |
| Oak & Maple vs Oak & Birch | 0.265 | 1.126 | 0.093 | 0.239 |
| **Paired vs All** |  |  |  |  |
| Oak & Maple vs All Trees | 0.263 | 1.262 | 0.112 | 0.060 |
| Oak & Birch vs All Trees | 0.268 | 1.146 | 0.113 | 0.242 |
| Birch & Maple vs All Trees | 0.517 | 2.585 | 0.205 | **0.004\*** |

